# Inhibition of the tuft cell/ILC2 axis reduces gastric tumor development in mice

**DOI:** 10.1101/2022.02.16.480779

**Authors:** Ryan N O’Keefe, Annalisa LE Carli, David Baloyan, Shoukat Afshar-Sterle, Moritz F Eissmann, Ashleigh R Poh, Cyril Seillet, Richard M Locksley, Matthias Ernst, Michael Buchert

## Abstract

Although gastric cancer is a leading cause of cancer-related deaths, systemic treatment strategies remain scarce. Here we explore a metabolite-triggered circuit between epithelial tuft cells and innate lymphoid type 2 cells (ILC2) that is evolutionarily optimized for intestinal remodeling in response to helminth infection. We demonstrate that tuft cell-derived interleukin 25 (IL25) acts as an alarmin on ILC2s to induce the release of IL13 as a growth factor for tuft cells, and propose that this model drives early metaplastic remodeling and gastric tumor formation. Genetic ablation of tuft cells, ILC2s or antibody-mediated neutralization of IL13 or IL25 reduces the growth of established tumors. Thus, the tuft cell/ILC2 axis provides an opportunity to therapeutically inhibit preneoplastic lesions and early-stage gastric cancer through repurposing of antibody-mediated therapies.

**One-Sentence Summary:** Tuft cells and type 2 innate lymphoid cells offer a new therapeutic target in gastric disease.

## Main Text

Gastric cancer is the 3^rd^ leading cause of cancer-related mortality worldwide, accounting for over a million new cases annually and close to 900,000 deaths (*1, 2*). Gastric cancer is predicted to increase in both incidence and mortality rates by approximately 40% by 2030 (*2, 3*), emphasising the need for greater understanding of molecular drivers of the disease. Intestinal metaplasia is a frequent non-malignant preneoplastic precursor to gastric cancer (*4*), and can be triggered by chronic gastritis as a consequence of persistent bacterial infection (*5*). The latter is characterised by epithelial tissue remodelling, loss of gastric acid-secreting parietal cells, and the expansion of metaplastic cells that resemble intestinal goblet cells (*6*). In addition to chronic gastric metaplasia, genetic, environmental and lifestyle factors such as smoking and alcohol consumption also increases the risk of gastric cancer (*7, 8*).

Tuft cells are a rare population of epithelial chemo-sensory cells that line the gastrointestinal and respiratory tract (*9, 10*), and can be identified by expression of Doublecortin-like kinase 1 (DCLK1) or Choline acetyltransferase (ChAT) in mouse and humans, respectively (*11*). In response to stimuli such as helminth infections, tuft cells secrete cytokines (e.g. interleukin (IL) 25), inflammatory mediators (e.g. eicosanoids), neurotransmitters (e.g. acetylcholine), and other signalling molecules that promote immune cell activation and tissue homeostasis (*10, 12–18*). However, emerging evidence suggests that tuft cells may also orchestrate early oncogenic processes, as evidenced by their rapid expansion and cancer stem cell-like properties in gastrointestinal pre-neoplastic lesions (*9, 19, 20*).

Tuft cell-derived IL25 promotes the activation of type 2 innate lymphoid cells (ILC2s), and the subsequent production of IL13 (*14–16*). ILC2s have come into focus within gastrointestinal tissues, for spearheading the immune defence against intestinal parasites and increasingly as a novel immune cell type regulating tumor immune responses (*21, 22*). Recent findings identified two major ILC2 resting and effector states (*23, 24*), which broadly correspond to natural (nILC2s) and inflammatory (iILC2s) (*25*). Tissue resident nILC2s are characterised by the expression of the IL33 receptor (ST2), and are involved in maintaining barrier homeostasis and repair (*26–28*). In contrast, iILC2s do not express ST2, and are recruited into mucosal tissues from the blood where they expand *in situ* in response to infection or IL25 signalling via the IL17RB receptor subunit (*26, 29, 30*).

The reciprocal tuft cell and ILC2 relationship is exemplified by their response to invading pathogens, where both populations are indispensable for the clearance of intestinal parasites and maintenance of tissue homeostasis (*31, 32*). Indeed, with elucidation of a feed-forward circuit between tuft cells and ILC2s in driving epithelial stem cell proliferation and tissue remodelling in the small intestine (*17*), we sought to examine a role for this processive circuit within mouse models of gastric disease and confirm its presence within human intestinal-type gastric cancer.

## Results

### Tuft cells and ILC2s are increased during Spasmolytic polypeptide-expressing metaplasia (SPEM), a precursor to gastric disease

Intestinal metaplasia is the leading risk factor for gastric cancer, while increased abundance of tuft cells and ILC2s have been reported in the gastric mucosa in response to *H. pylori* infection (*33, 34*) and metaplasia (*35–38*). The latter is experimentally replicated following a 3-day exposure to high dose tamoxifen (HDTmx; 250mg/kg) (*39*) and results in the loss of parietal cells and H^+^/K^+^ATPase expression, with a concomitant trans-differentiation of gastric chief cells into a spasmolytic polypeptide-expressing metaplasia (SPEM) characterized by TFF2 expression (Fig. S1B) (*39*). To assess the contribution of the tuft cells/ILC2 axis in gastric metaplasia, we quantified the abundance of these cells 2 days after HDTmx administration, and observed an increased proportion of SiglecF^+^CD24^+^EpCAM^+^ tuft cells and KLRG1^+^ST2^+/-^CD90.2^+^ ILC2s compared to the vehicle-treated controls (Fig. S1C/D, S2A/B). Strikingly, we found that the increase in total ILC2s was almost entirely due to an increase in the IL25-responsive iILC2 subpopulation (Fig. S1E), whereas the number of IL33-responsive nILC2s remained unchanged (Fig. S1F).

Next, we utilised *BAC(Dclk1^CreERT2^);Rosa26^DTA/+^* tuft cell *deleter* (TC*^i-DTA^*) mice to establish a functional role for tuft cells during gastric disease. This strain affords the ablation of tuft cells in response to the activation of a diphtheria toxin A (*DTA*) allele by the tuft cell-specific and tamoxifen-inducible *Dclk1^CreERT2^* driver (*40*). We treated TC*^i-DTA^* and *Dclk1^+/+^;Rosa26^DTA^* (TC*^WT^*) control mice with low dose tamoxifen (LDTmx; 50mg/kg, Fig. 1A) to solely activate the cre recombinase, or with HDTmx to induce SPEM in the absence of tuft cells (Fig. 1B). Interestingly, TC*^i-DTA^* mice treated with HDTmx were partially protected from the loss of parietal cells and SPEM (Fig. 1C). Strikingly, both tamoxifen treatment regimens induced a near complete ablation of tuft cells within the gastric epithelium (Fig. 1D), alongside a significant reduction of ILC2s in TC*^i-DTA^* mice (Fig. 1E). Interestingly, while iILC2S were unaffected by tuft cell depletion following LDTmx treatment (Fig. 1G), HDTmx significantly reduced both nILC2 and iILC2 populations (Fig. 1F/G). Collectively, these results suggest that tuft cells may regulate the presence of iILC2 in the gastric mucosa, while SPEM may induce abundance of iILC2 in a tuft cell-dependent manner.

**Fig. 1.**
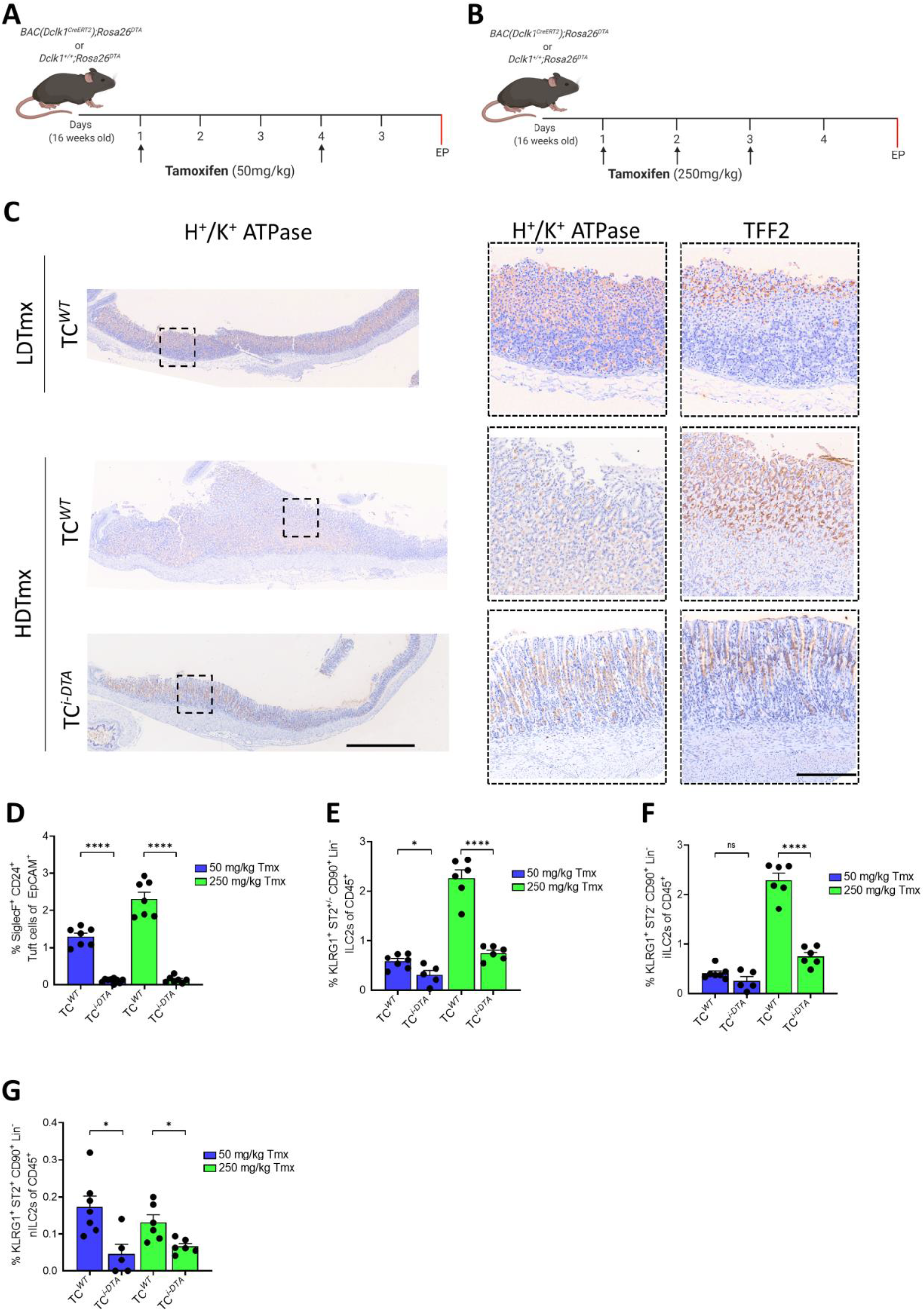
Ablation of tuft cells results in reduced ILC2s in both WT and HDTmx treated mice. (**A**) Experimental schematic for tuft cell ablation. 16-week old *Dclk1^+/+^; Rosa26^DTA^* (TC*^WT^*) and inducible *BAC(Dclk1^CreERT2^);Rosa26^DTA/+^* (TC*^i-DTA^*) mice administered 2 injections of LDTmx (50 mg/kg) to induce tuft cell ablation (in *Dclk1^CreERT2^;Rosa26^DTA^* mice). EP = endpoint. (**B**) Experimental schematic for SPEM induction and tuft cell ablation. 16-week-old TC*^WT^* and TC*^i-DTA^* mice were treated with HDTmx (250 mg/kg) once daily for 3 consecutive days to induce gastric spasmolytic polypeptide-expressing metaplasia (SPEM) and tuft cell ablation (in *Dclk1^CreERT2^;Rosa26^DTA^* mice). EP = endpoint. (**C**) Representative immunohistochemical (IHC) images of stomachs from TC*^WT^* and TC*^i-DTA^* mice treated as described in Fig. 1A. Scale bar = 1 mm and 300 μm (dotted insets). (**D**) Flow-cytometry quantification of SiglecF^+^CD24^+^EpCAM^+^ tuft cells from stomachs of TC*^WT^* and TC*^i-DTA^* mice treated with LDTmx or HDTmx. (**E**) Flow-cytometry quantification of KLRG1^+^ST2^+/-^CD90.2^+^Lineage^-^CD45^+^ ILC2s in stomachs of TC*^WT^* and TC*^i-DTA^* mice treated with LDTmx or HDTmx. (**F**) Flow-cytometry quantification of KLRG1^+^ST2^-^CD90.2^+^Lineage^-^CD45^+^ ILC2s in stomachs of TC*^WT^* and *TC^i-DTA^* mice treated with LDTmx or HDTmx. (**G**) Flow-cytometry quantification of KLRG1^+^ST2^+^CD90.2^+^Lineage^-^CD45^+^ ILC2s in stomachs of TC*^WT^* and TC*^i-DTA^* mice treated with LDTmx or HDTmx. Data represents mean ± SEM, p values from ANOVA * p < 0.05, **** p< 0.0001, n.s - not significant. Each symbol represents an individual mouse.

### Tuft cells and ILC2s are increased in during gastric tumor development

Given the strong predisposition of cancer development associated with SPEM, we next dissected the role of tuft cells and ILC2s during early gastric tumor development in the *Gp130*^F/F^ mouse model, which spontaneously develops intestinal-type gastric adenomas from 4 weeks of age (Fig. S3A) (*41*). Compared to the normal mucosa of control WT mice (*Gp130^+/+^*), adenomas of *Gp130*^F/F^ mice demonstrated an increased proportion of DCLK1^+^ and SiglecF^+^CD24^+^EpCAM^+^ tuft cells (Fig. S3C-D), which coincided with an increased abundance of iILC2s (Fig. S3E-3G).

To confirm a tumor-promoting role for tuft cells in gastric cancer, we created an inducible *Gp130^F/F^;BAC(Dclk1^CreERT2^);Rosa26^DTA/+^* (referred to as *Gp130^F/F^*;TC*^i-DTA^*) compound mutant mouse. Following LDTmx administration, *Gp130^F/F^*;TC*^i-DTA^* mice developed smaller tumors compared to *Gp130^F/F^*;TC*^WT^* mice (Fig. 2A/B/C). In support of the relationship between tuft cells and ILC2s in tumor formation, ILC2-deficient *Gp130^F/F^*;R5-IL5^dtTomato-IRES-Cre^*;Rosa26*^DTA/+^ (referred to as *Gp130^F/F^*;ILC2*^c-DTA^*) exhibited a reduced tumor burden compared to *Gp130^F/F^*;ILC2*^WT^* mice (Fig. 2B-2D), which coincided with significantly fewer tuft cells (Fig. 2E). Conversely, we observed a significant reduction in total ILC2s within the stomachs of *Gp130^F/F^;TC^i-DTA^* mice (Fig. 2F), which was attributed primarily to reduced iILC2s (Fig. 2G, 2H). Together our findings demonstrate a functional role for tuft cells and ILC2s in promoting early gastric tumor development in mice, expanding the interdependency between tuft cells and ILC2 observed in situations of SPEM to that of adenoma formation and tumor development.

**Fig. 2.**
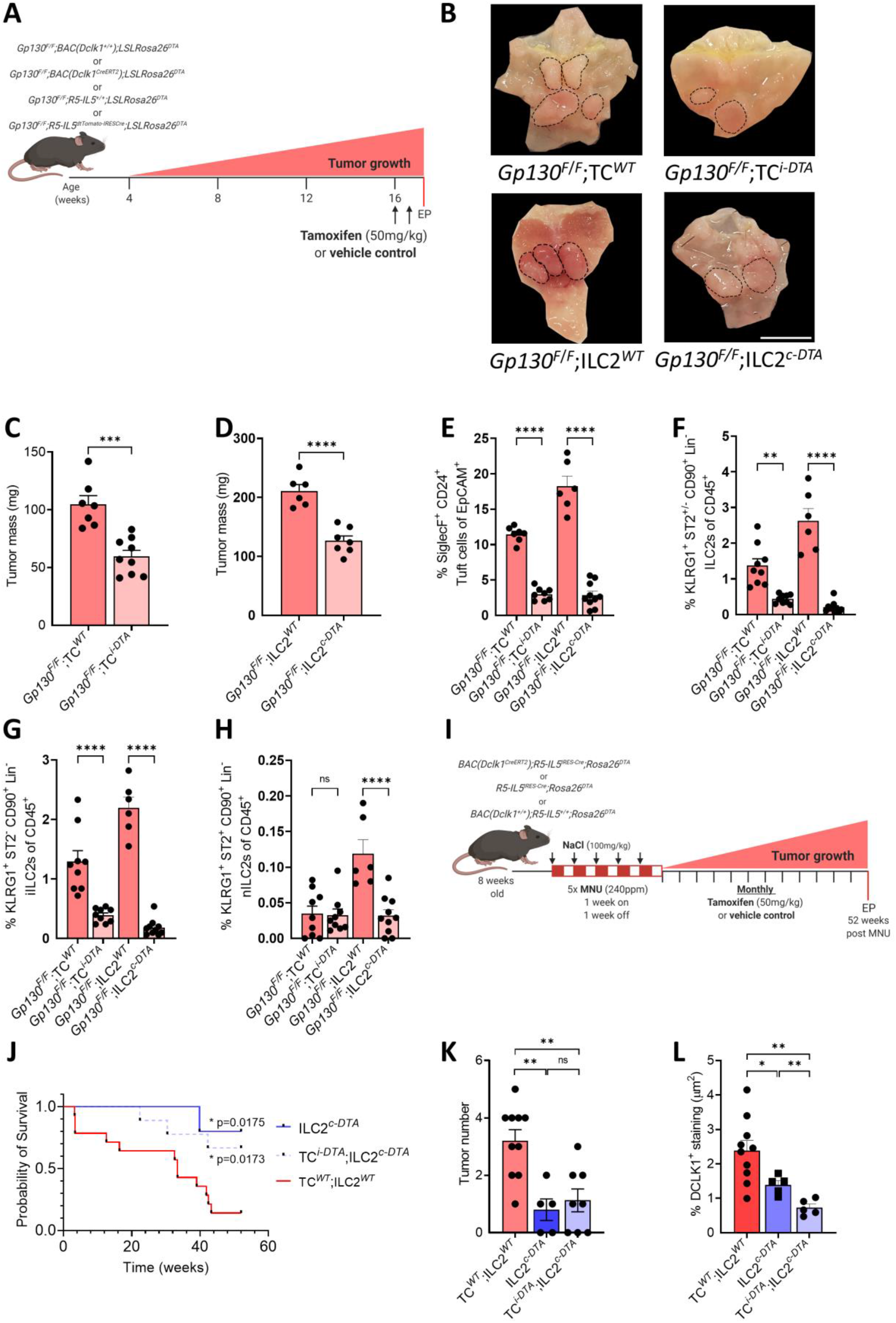
Genetic ablation of tuft cell reduces gastric adenoma growth. (**A**) Schematic outlining the genetic mouse models used in Fig. 2 to ablate tuft cells and ILC2s. Tuft cell ablation was achieved in *Gp130^F/F^;BAC(Dclk1^CreERT2^);Rosa26^DTA/+^* (TC*^i-DTA^*) mice and compared against *Gp130^F/F^;Dclk1^+/+^;Rosa26^DTA^* (*Gp130^F/F^*;TC*^WT^*) littermate controls. Constitutive ILC2 ablation in *Gp130^F/F^;R5-IL5^dtTomato-IRESCre^;LSLRosa26^DTA^* (ILC2*^c-DTA^*) mice was compared against *Gp130^F/F^;R5-IL5^+/+^;LSLRosa26^DTA^* (*Gp130^F/F^*;ILC2*^WT^*) littermate controls. All cohorts were administered 2 injections of tamoxifen to induce tuft cell ablation at 16-weeks of age. EP = endpoint. (**B**) Representative stomach images of *Gp130^F/F^*;ILC2*^WT^*, *Gp130^F/F^*;TC*^WT^*, *Gp130^F/F^*;TC*^i-DTA^* and *Gp130^F/F^*;ILC2*^c-DTA^* mice treated as described in Fig. 2A Black dotted circles indicate tumors, scale bar = 8 mm. (**C**) Tumor mass of 17-week-old *Gp130^F/F^* mice following genetic tuft cell ablation compared to vehicle control treated *Gp130^F/F^*;TC*^WT^*. (**D**) Tumor mass of 17-week-old ILC2-deficient *Gp130^F/F^*;ILC2*^c-DTA^* mice compared to age matched *Gp130^F/F^*;ILC2*^WT^* mice. (**E**) Flow-cytometry quantification of SiglecF^+^ CD24^+^ EpCAM^+^ tuft cells in tumors of the indicated genotypes treated as described in Fig. 2A. (**F**) Flow-cytometry quantification of ILC2s as KLRG1^+^ST2^+/-^CD90.2^+^Lineage^-^CD45^+^ in tumors of the indicated genotypes treated as described in Fig. 2A. (**G-H**) Flow-cytometry quantification of KLRG1^+^ST2^-^CD90.2^+^Lineage^-^CD45^+^ iILC2s and KLRG1^+^ST2^+^CD90.2^+^Lineage^-^CD45^+^ nILC2s in tumors of the indicated genotypes treated as described in Fig. 2A. (**I**) Experimental schematic outlining the genetic mouse model of MNU/NaCl-induced gastric cancer. *Dclk1^CreERT2^;R5-IL5^dtTomato-IRESCre^;Rosa26^DTA^* (TC*^i-DTA^*;ILC2*^c-DTA^*), ILC2*^c-DTA^* and *DclkG1^+/+^;R5-IL5^+/+^;Rosa26^DTA^* (ILC2*^WT^*;TC*^WT^*) mice were treated with MNU for 5 alternating weeks, then were aged for 52 weeks with monthly tamoxifen injections to ablate tuft cells. EP = endpoint. (**J**) Kaplan-Meier survival analysis of MNU treated mice, ILC2*^c-DTA^* * p=0.0175 (cp to ILC2*^WT^*;TC*^WT^*), TC*^i-DTA^*;ILC2*^c-DTA^* * p=0.0173 (cp to ILC2*^WT^*;TC*^WT^* Mantel-Cox test). (**K**) Quantification of tumor numbers in MNU-treated mice as described in Fig. 2I. (**L**) IHC quantification of DCLK1^+^ cells in MNU treated TC*^WT^*;ILC2*^WT^*, TC*^i-DTA^*;ILC2*^c-DTA^* and ILC2*^c-DTA^* mice. Data represents mean ± SEM, p values from Student’s t-test or ANOVA * p < 0.05, ** p < 0.01, *** p< 0.001, **** p< 0.0001, n.s - not significant. Each symbol represents an individual mouse.

To confirm an involvement of tuft cells and ILC2s in sporadically occurring gastric cancer, we next induced gastric adenocarcinomas with the carcinogen N-Methyl-N-Nitrosourea (MNU) in WT mice, ILC2-deficient mice (*R5-IL5^dtTomato-IRES-Cre^;LSL-Rosa26^DTA^;* referred to as ILC2*^c-DTA^* (*42*)), and ILC2*^c-DTA^* mice with conditional tuft cell ablation (referred to as TC*^i-DTA^*;ILC2*^c-DTA^*)(Fig. 2I). We observed extended survival of ILC2*^c-DTA^* mice compared to their WT littermates (Fig. 2J), regardless tuft cell ablation. Additionally, ILC2*^c-DTA^* and TC*^i-DTA^*;ILC2*^c-DTA^* mice both developed significantly fewer gastric adenocarcinomas than control WT mice (Fig. 2K). Consistent with the observation that tuft cell ablation reduces gastric cancer burden in MNU-treated mice (*43*), we identified fewer tuft cells in both TC*^i-DTA^*;ILC2*^c-DTA^* and ILC2*^c-DTA^* cohorts (Fig. 2L). Collectively, our data identifies tuft cells and ILC2s as key drivers of gastric adenocarcinoma formation.

### Tuft cell/ILC2 signalling is primarily through IL13 and IL2

The increased abundance of tuft cells and ILC2s in the gastric mucosa and adenomas of *Gp130^F/F^* mice is reminiscent of an intestinal, canonical type 2 anti-helminth immune response, where tuft cells secrete IL25 upon detection of helminth metabolites (*17, 31*) and results in the expansion of intestinal iILC2s. In turn, these cells secrete IL13 which promotes the differentiation of intestinal progenitor cells towards goblet cell and tuft cell lineages (*14–16*). Importantly, this process is impaired in the absence of IL25 or IL13, delaying parasite clearance (*14–16*). Surprisingly, we observed a similar correlation between elevated expression of the tuft cell and ILC2 genes *Dclk1* and *Gata3* and the cytokines *Il25* and *Il13* following induction of SPEM in WT mice (Fig. S4A). We functionally confirmed this correlation in the low and high dose tamoxifen experiments in our TC*^i-DTA^* mice, which showed significantly decreased expression of the prototypical ILC2 genes *Gata3* and *Il13* (Fig. S4B). Conversely, tumors of *Gp130^F/F^* mice also expressed more *Dclk1, Il25, Gata3* and *Il13* than the normal stomach tissue of wildtype *Gp130^+/+^* mice (Fig. S4C). Finally, tuft cell-deficient *Gp130^F/F^;*TC*^i-DTA^* mice demonstrated reduced expression of *Dclk1, Il25, Gata3* and *Il13* (Fig. S4D). These observations were reproduced in MNU-treated mice, where *Dclk1, Il25, Gata3* and *Il13* expression was reduced in adenocarcinomas across TC*^i-DTA^*;ILC2*^c-DTA^* and ILC2*^c-DTA^* cohorts (Fig. S4E).

### Pharmacologic inhibition of the tuft cell / ILC2 signaling loop reduces gastric tumor growth

Based on the above, we next explored whether inhibition of IL25 or IL13 signalling would limit gastric tumor growth and treated tumor-bearing *Gp130^F/F^* mice with either α-IL13 or α-IL25 neutralizing antibodies (Fig. 3A). Compared to *Gp130^F/F^* mice from the IgG-treated control cohorts, both α-IL13 and α-IL25 treatment reduced tumor growth (Fig. 3B-3D). This also coincided with a reduced proportion of tuft cells (Fig. 3E, 3F), iILC2s (Fig. 3G, 3H) and their respective *Dclk1, Il25, Gata3* and *IL13* genes within the gastric adenomas and the surrounding mucosa (Fig. S4F, S4G). Because FACs purified tuft cells from the stomachs of *Gp130^+/+^* and *Gp130^F/F^* mice expressed significantly more *Dclk1, Il25* and *Il13Ra1* (encoding the IL13 receptor) than their EpCAM^+^ epithelial counterparts (Fig. S4H), we conclude that gastric tuft cells, like their intestinal counterparts, are the primary source of IL13-dependent IL25 production. Interestingly, we observed that *Il13* was exclusively expressed by ILC2s from *Gp130^F/F^* gastric adenomas (Fig. S4I), while expression of *Gata3* and the *Il25* receptor component *Il17rb* was dramatically increased in ILC2s compared to CD45^+^ cells isolated from wildtype *Gp130^+/+^* and *Gp130^F/F^* mice (Fig. S4I). These findings suggest that under homeostatic conditions the relatively low levels of tuft cell derived IL25 within the gastric epithelium may be insufficient to activate the ILC2s residing in the gastric mucosa, reminiscent of the small proportion of IL13 expressing ILC2s in the intestine under homeostatic conditions (*15*). Collectively, our data point to a tuft cell/ILC2 circuit that is controlled by reciprocal production of IL25/IL13, and possibly amplified by a feed forward loop through the IL25-response of IL17rb expressing tuft cells (*17*), as a phylogenetically conserved defence mechanism against helminth infections and coerced in neoplastic situations. Importantly, this circuit can be therapeutically exploited to suppress growth of gastric malignancies with clinically relevant anti-cytokine monoclonal antibodies.

**Fig. 3.**
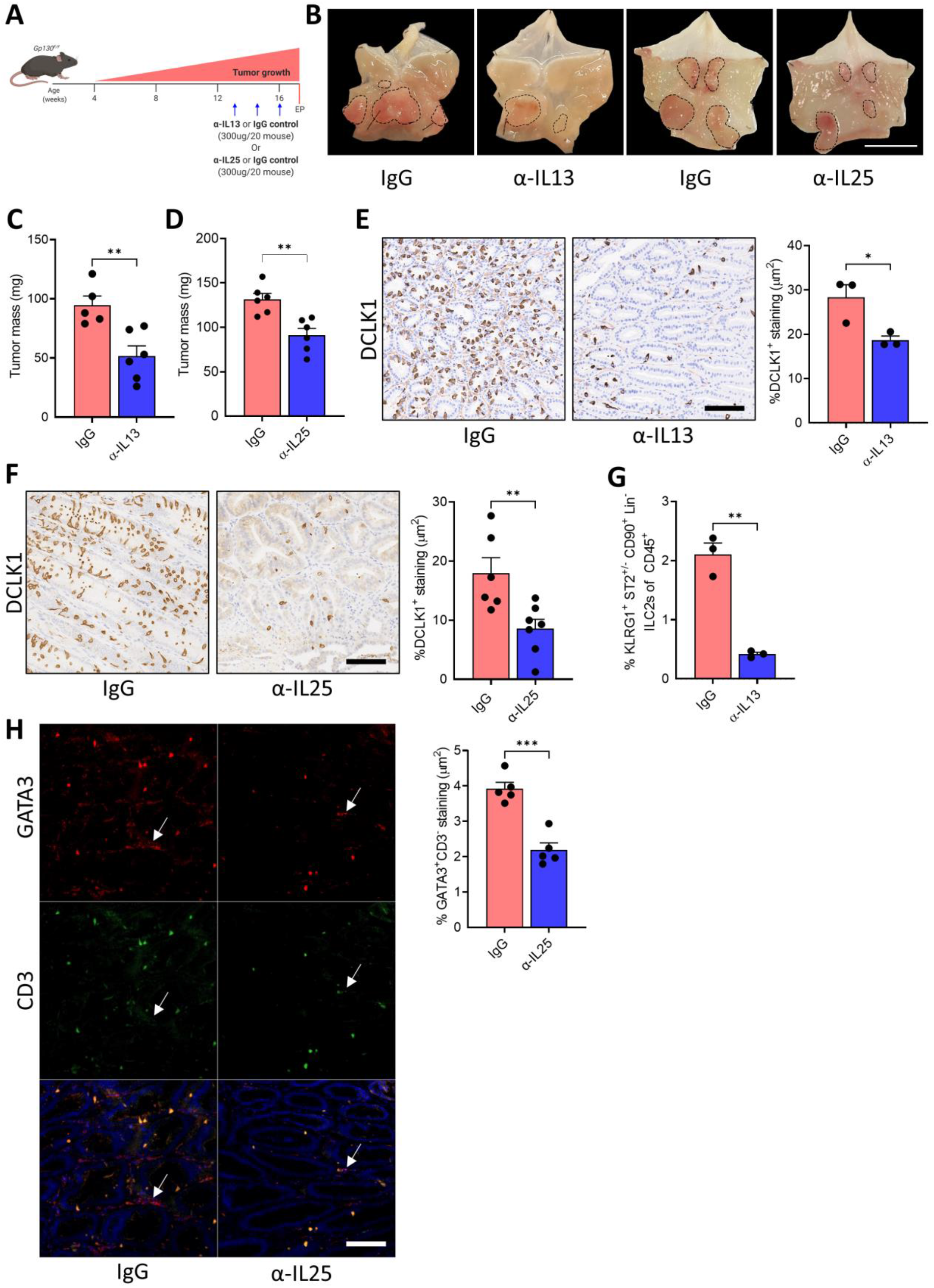
Pharmacologic inhibition of the tuft cell / ILC2 signaling loop reduces gastric tumor growth. (**A**) Schematic outlining the antibody-mediated blockade of IL13 or IL25. 13-week-old mice were treated with either α-IL13, α-IL25 or a matched IgG isotype control (300 μg/20g mouse, once weekly for 3 weeks). Mice were culled one week after the third injection. EP = endpoint. (**B**) Representative images of *Gp130^F/F^* stomachs treated as described in Fig. 3A. Dotted circles indicate tumors, scale bar = 8 mm. (**C**) Tumor mass in *Gp130^F/F^* mice following treatment with α-IL13 or a matched IgG isotype control. (**D**) Tumor mass of *Gp130^F/F^* mice following treatment with α-IL25 or a matched IgG isotype control. (**E**) IHC staining and quantification of DCLK1^+^tuft cells in α-IL13 and IgG treated *Gp130^F/F^* mice, scale bar = 300 μm. (**F**) IHC staining quantification of DCLK1^+^ tuft cells in α-IL25 and IgG treated *Gp130^F/F^* mice, scale bar = 300 μm. (**G**) Flow-cytometry quantification of KLRG1^+^ST2^+/-^CD90.2^+^Lineage^-^ CD45^-^ ILC2s in tumors of *Gp130^F/F^* mice treated with α-IL13 or a matched IgG isotype control. (**H**) Immunofluorescence (IF) staining and quantification of Gata3^+^CD3^-^ ILC2s in tumors of *Gp130^F/F^* mice treated with α-IL25 or a matched IgG control. Arrows indicate ILC2s, scale bar = 100 μm. Data represents mean ± SEM, p values from Student’s t-test * p <0.05, ** p < 0.01, *** p< 0.001. Each symbol represents an individual mouse.

### Tuft cells and ILC2 are involved in human gastric cancer

We next sought to determine whether the tuft cell/ILC2 signalling circuit could predict clinical outcome in patients. We quantified the abundance of tuft cells and ILC2s in tumor microarrays of intestinal-type gastric cancer patients by staining for choline O-acetyltransferase (ChAT) and GATA3^+^CD3^-^ cells, respectively (*43, 44*). We observed co-existence of tuft cells and ILC2s in 40% of all tumor specimens (n=67) (Fig. 4A, 4B). Given that tuft cells are the primary source of epithelial cell-derived IL25 in the gastrointestinal and respiratory tract (*15, 45*), we next interrogated survival data for intestinal-type gastric cancer patients against the median expression level of a manually-curated human tuft cell gene signature (*ChAT, AVIL, IL25*) (*46*). We found that patients that highly expressed high levels of *ChAT/AVIL/IL25* exhibited significantly poorer overall survival compared to patients that expressed low levels of the tuft cell signature (Fig. 4C). Likewise, patients with high expression of ILC2 marker genes (*GATA3* and *IL13*) had poorer overall survival compared to patients with low expression (Fig. 4C). Significantly, the correlation of expression signatures for tuft cells, ILC2s and survival was unique to intestinal-type gastric cancer, while there was no correlation with patient outcome within diffuse-type gastric cancer (Fig. 4D). We therefore conclude that the functional involvement of the tuft cell/ILC2 circuit in preclinical mouse models is highly likely to functionally contribute to the progression of intestinal-type gastric cancers in humans.

**Fig. 4.**
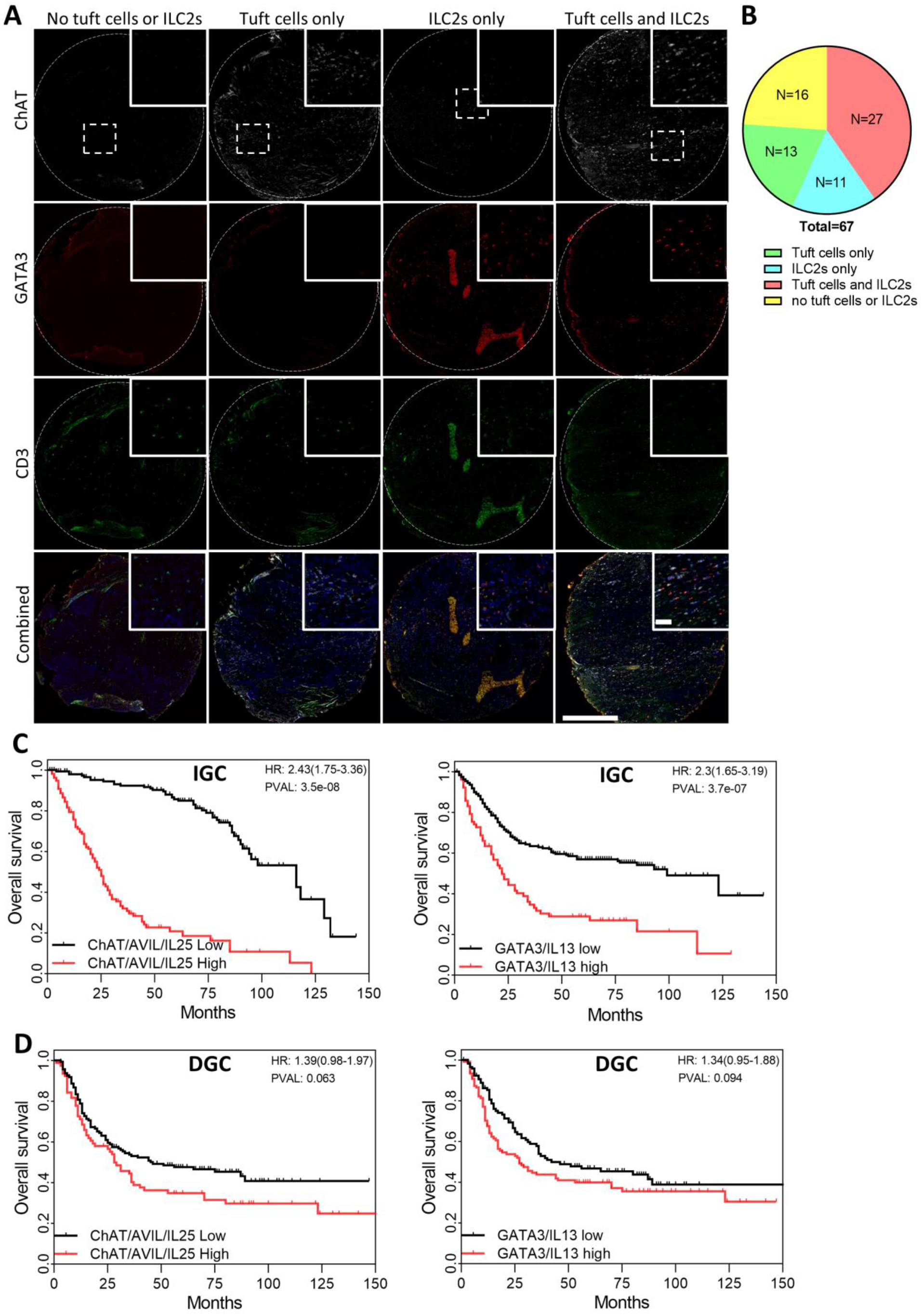
Tuft cells and ILC2 are involved in human gastric cancer. (**A**) Representative IF stains for ChAT^+^ tuft cells and CD3^-^GATA3^+^ ILC2s in human intestinal-type gastric cancer tissue array. Scale bar = 500 μm for low magnification and =100 μm for high magnification (dotted insets). (**B**) Quantification of tuft cells and/or ILC2s in human intestinal-type gastric cancer tumor micro-arrays. (**C-D**) Kaplan-Meier survival analysis for intestinal-type (IGC) and diffuse-type (DGC) gastric cancer patients segregated at the median level of gene expression for tuft cell (*ChAT, AVIL* and *IL25*) and ILC2 (*GATA3* and *IL13*) gene signatures.

## Discussion

The susceptibility of cancer promotion to interference with anti-cytokine (signaling) therapy provides novel and exciting therapeutic opportunities to target cancer cell intrinsic hallmarks as well as shape the stromal and immune response of the tumor environment. Here we describe a novel regulatory circuit between epithelial tuft cells and ILC2s connected through reciprocal production of, and response to, IL25 and IL13. Although cytokine responsiveness of ILC2s is contextual, IL33 and IL25 remain the major driver (*47*), with prominent roles for IL25 during skin allergies (*48*), pulmonary fibrosis (*49*) and helminth defense (*50*). A regulatory tuft cells/ILC2 circuit had previously been demonstrated to drive the clearance of invading intestinal parasites in murine models (*16, 51, 52*). Indeed, the expansion of the tuft cell and ILC2 populations required to overcome parasite infections are also cellular hallmarks of epithelial metaplasia in the stomach. Meanwhile, tuft cell and ILC2 numbers have separately been found to increase during gastric colonization with *H. pylori* infection, and chemically induced gastric metaplasia (*36–38*), precursors of chronic gastritis and associated progression to gastric cancer (*33, 34*). In agreement with our findings, genetic or antibody-mediated depletion of ILC2s has been demonstrated to protect against chemically induced gastric metaplasia (*37, 38*), although this was shown in the context of IL13 and IL33 signaling, while IL25 was not investigated (*37*).

It is currently unclear how IL25-secreting tuft cells initially expand in the gastric mucosa. It has previously been reported that the expansion of gastric tuft cells during SPEM is dependent on gastrin released by endocrine cells, however subsequent lineage tracing experiments have failed to identify the gastric stem cell responsible for producing tuft cells during SPEM (*36*). While in the small intestine IL13 signals to stem/progenitor cells promotes hyperplasia of IL25^+^ tuft cells, in the stomach IL13 receptor expression has been reported in both chief cells and metaplastic SPEM cells (*53*). Given that we observe increased levels of *de novo* expression of *Il13ra1* in FACs isolated tuft cells from *Gp130^F/F^* mice compared to wildtype, we speculate that tuft cell expansion during metaplasia and adenoma formation is driven by one or both of these cell populations. While the tuft cell/ILC2 circuit is maintained by IL25 and IL13 cytokine signaling, it remains unclear how this feed-forward loop is initiated. Recently, cysteine leukotrienes secreted by tuft cells, have been shown to be required for full ILC2 activation downstream of IL25 in the promotion of helminth clearance in the intestine (*51*). While it is tempting to speculate that leukotrienes may be complementing IL25 signaling during gastric disease development, our data also demonstrate that blocking IL25 alone was sufficient to impair iILC2 expansion and tumor development. Interestingly, in a comparable mouse model where genetic ablation of either *Il33* or *Il13* expression prevented experimentally induced gastric metaplasia, IL33 is secreted by either metaplasia-associated macrophages or through epithelial cell damage and induced the release of IL13 among other Th2 cytokines, predominantly from ILC2s (*37, 53*). Thus, it is feasible that during parietal cell loss, ILC2s are initially activated by either epithelial cell damage or macrophage derived IL33, which in turn induces tuft cell expansion by IL13 signaling in gastric chief and/or metaplastic SPEM cells. Once the tuft cell population has sufficiently expanded, IL25 becomes the dominant driver for the expansion and excessive activation of the IL25-responsive population of tissue-resident mucosal ILC2s.

Our study demonstrates that the tuft cell/ILC2 circuit phylogenetically optimized as a repair mechanism of intestinal epithelium during helminth infection becomes hijacked to promote the progression of gastric cancer both at early metaplastic and adenomatous, as well as at later carcinoma stages. This circuit is maintained by complementary IL25 and IL13 signalling between the two cell types. Importantly, both genetic interference of the circuit through ablation of either tuft cells or ILC2s, or therapeutic suppression of either IL13 or IL25 signalling confers a profound reduction of disease burden at the earliest stages (i.e. gastric metaplasia) as well as with established metaplastic lesions (i.e. gastric adenomas and adenocarcinomas). We predict that our functional insights in preclinical models will translate to humans, as tuft cell and ILC2 expression “signatures” are associated with worse survival in patients suffering from intestinal-type gastric cancer, where roughly one half of all tumors show enrichment of both tuft cells and ILC2s. A swift clinical translation of our discovery is supported by the availability of α-IL13 monoclonal antibodies that are currently optimized for the treatment of severe asthma (*54*). This opens the possibility of developing new treatment strategies and modalities not only for gastric cancer, but also for companion diagnostics for early detection and patient stratification.

## Acknowledgments

We are indebted to David Williams (Department of Pathology, Austin Hospital) for providing the gastric cancer tissue arrays. We thank the members of the Cancer and Inflammation Program at the Olivia Newton-John Cancer Research Institute for helpful discussions and comments. Experimental schematics were created with BioRender.com.

## Funding

National Health and Medical Research Council of Australia (NHMRC) Principal Research Fellowship 1079257 (ME)

NHMRC Program Grant 1092788 (ME)

NHMRC Investigator Grant 1173814 (ME)

NHMRC Project Grant 1143020 (MB)

La Trobe RFA Understanding Disease Grant (MB)

Operational Infrastructure Support Program, Victorian Government, Australia (MB)

La Trobe University Graduate Research Scholarship (RNO)

NHMRC Peter Doherty Early Career Fellowship GNT1166447 (ARP)

NHMRC Project Grant 1143030 (RML)

National Institute of Health AI026918 (RML)

Howard Hughes Medical Institute (RML)

The Sandler Asthma Basic Research Center (RML)

Cancer Council Victoria’s Grant-in-Aid APP1160708 (MFE)

Victorian Cancer Agency Mid-Career Research Fellowship MCRF20018 (MFE)

## Author contributions

Conceptualization: MB, ME, RNO, RML

Methodology: RNO, MB, RML, CS, ARP

Investigation: RNO, MB, ALC, SAS, ARP

Resources: MB, ME, RML, DB, MFE

Visualization: RNO, MB, ARP

Funding acquisition: MB, ME

Project administration: MB, ME

Supervision: MB, ME

Writing – original draft: RNO, MB, ALC

Writing – review & editing: RNO, MB, ME, RML, CS, ALC, MFE, ARP

## Data and materials availability

All data are available in the main text or the supplementary materials. Source data and reagents used can be made available upon reasonable request.

## Supplementary Materials

Materials and Methods

Figs. S1 to S4

Tables S1 to S3

