## Additional data for "Inhibition of the tuft cell/ILC2 axis reduces gastric tumor development in mice"

**This PDF file includes:**

Materials and Methods
Figs. S1 to S4
Tables S1 to S3

### Materials and Methods

***Study Approval:*** All animal studies were conducted in accordance with all relevant ethical regulations for animal testing and research including the Australian code for the care and use of animals for scientific purposes. All animal studies were approved by the Animal Ethics Committee of Austin Health (A2019_05602, A2015_05289) or La Trobe University (AEC 17-73). We have complied with all relevant ethical regulations for work with human participants. Collection and usage of human gastric cancer tissues was approved by the Austin Health ethics committee (HREC/15/Austin/359) and informed consent was obtained from all subjects.

***Animal Models:*** All mice were bred and maintained under specific pathogen-free conditions in the research facility of the La Trobe University or Austin Health. All strains were maintained on a 12-hour light/dark cycle at constant temperature. Co-housed, age- and gender-matched littermates were utilized for all experiments. All interventions were performed during the light cycle on both male and female mice. All animals had free access to water and food (standard chow). The inducible $BAC(Dclk1^{CreERT2})$ strain has been previously reported (*37*) and was crossed with $LSL\text{-}Rosa26^{DTA}$ model to generate a mouse model of tuft cell ablation and with $Gp130^{Y757F}$ ($Gp130^{F/F}$) , a murine model of gastric cancer (*38*), to generate $BAC(Dclk1^{CreERT2});Gp130^{F/F};LSL\text{-}Rosa26^{DTA}$, a murine model of gastric cancer with inducible tuft cell ablation. The $R5\text{-}IL5^{dtTomato\text{-}IRESCre};LSLRosa26^{DTA}$ model of constitutive ILC2 depletion has previously been reported (*39*) and was crossed with $Gp130^{F/F}$ or $BAC(Dclk1^{CreERT2})$ mice to generate the $Gp130^{F/F};R5\text{-}IL5^{dtTomato\text{-}IRESCre};LSLRosa26^{DTA}$ strain, a murine model of gastric cancer with constitutive ILC2 depletion, or $BAC(Dclk1^{CreERT2});R5\text{-}IL5^{dtTomato\text{-}IRESCre};LSLRosa26^{DTA}$ mice, respectively, a mouse model lacking ILC2s with optional (tmx-inducible) tuft cell ablation. The control cohorts for tuft cell ablated mice, were comprised of $Dclk1^{+/+};Rosa26^{DTA}$, or $Gp130^{F/F};Dclk1^{+/+};Rosa26^{DTA}$ age matched littermates. Control cohorts of ILC2 depleted mice were comprised of $R5\text{-}IL5^{+/+};LSLRosa26^{DTA}$ or $Gp130^{F/F};R5\text{-}IL5^{+/+};LSLRosa26^{DTA}$ mice age matched littermates.

***Tissue collection:*** Stomachs were removed, cut along the greater curvature and flushed with cold PBS to remove contents. Stomachs were pinned out, with tumors being excised using curved scissors, taking care to avoid the mucosa. Tumors and stomach tissue were either fixed in 10 % neutral buffered formalin (NFB) overnight at room temperature for histological analysis or snap frozen on dry ice and stored at -80°C for molecular analysis.

***Tamoxifen treatment:*** Low dose tamoxifen (LDTmx, Sigma, T5648) was prepared at a concentration of 10 mg/ml in 10 % ethanol and sterile sunflower oil. Mice were administered two doses of tamoxifen at 10 mg/kg via i.p. three days apart. Two days after the last tamoxifen dose mice were euthanized via $CO_2$ asphyxiation and their stomachs were collected.
For the high dose tamoxifen (HDTmx) treatment, tamoxifen was prepared at a concentration of 250 mg/ml in 10 % ethanol and sterile sunflower oil. Mice were administered tamoxifen at 250 mg/kg via i.p. once a day for three days. Two days after the last tamoxifen dose mice were euthanized via $CO_2$ asphyxiation and their stomachs were collected. Vehicle-treated control mice were administered 10% ethanol in sunflower oil.

***MNU treatment:*** 8-week-old mice were treated with a regimen of MNU (240 ppm) in the drinking water (1 week ON and 1 week OFF for 10 consecutive weeks). At the beginning of each MNU treatment week, mice were also administered 100 mg/kg of NaCl by oral gavage to increase the incidence of gastric tumor development (*52*). 52 weeks after the last MNU treatment mice were euthanized via $CO_2$ asphyxiation and stomachs were collected and tumor numbers assessed.

***α-IL25 and α-IL13 treatment:*** 13-week-old *Gp130*$^{F/F}$ mice were given 1x weekly injection for 3 weeks of either α-IL25 (R&D Systems, MAB13992), α-IL13 (R&D Systems, MAB413) or IgG control (R&D Systems, MAB006 and MAB004) (at 300 μg/mouse). 1 week after the last injection, mice were euthanized via $CO_2$ asphyxiation. Stomachs were collected and tumors were excised and weighed.

***TCGA dataset analysis:*** Using the online KM plotter tool (https://www.kmplot.com/), we interrogated the patient survival against the median expression level of tuft cell related gene (*ChAT, AVIL, IL25*), and ILC2 related genes (*GATA3 and IL13*) expression within following datasets; GSE14210, GSE15459, GSE22377, GSE29272, GSE51105 and GSE62254. We categorised intestinal-type gastric cancer patients and diffuse-type gastric cancer patients into groups with either high or low gene expression quartiles.

***RNA extraction and RT-PCR analysis:*** RNA extraction from whole tissue was performed using the RNeasy Mini Kit (QIAGEN, 74106), and cDNA was generated using the High-Capacity cDNA Reverse Transcription Kit (Applied Biosystems, 4368813) in accordance with the manufacturer's instructions. RNA extraction from sorted cells was performed using the RN-easy Micro Plus kit (Qiagen), cDNA was generated with the SuperScript™ IV First-Strand Synthesis System (ThermoFisher) in accordance with the manufacturer's instructions. Quantitative RT-PCR analysis was performed using the SensiMix SYBR Hi-ROX Kit (Bioline, QT605-20) in duplicates (technical replicates) using the Viia7 Real-Time PCR System (Life Technologies). Samples were exposed to an initial denaturation step of 95°C/10min, followed by 40 cycles of amplification (95°C for 15s, 60°C/1min). *18S* or *Gapdh* were used as house-keeping genes, with fold changes in gene expression being calculated using the 2-ΔΔCT method. Primers used are outlined in Table 1.

***Immunohistochemistry and quantification:*** Following fixation in 10% NFB, tissue was embedded in paraffin and cut into 10 mm thick sections. These sections underwent dewaxing and tissue hydration via incubation in xylene followed by gradient ethanol washes. Antigen retrieval was performed with citrate buffer heated in a microwave pressure cooker (pH 6 for 15 minutes), followed by blocking in 10 % (v/v) normal goat serum for 1 hour at room temperature. Primary antibodies as outlined in Table S2, were diluted in 10 % (v/v) normal goat serum and incubated overnight at 4$^{o}$C in a humidified chamber. Secondary HRP antibodies used were polyclonal rabbit anti-goat (Dako; P0449), polyclonal goat anti-rabbit (Dako; P0448), polyclonal goat anti-mouse (Dako; P0447), polyclonal goat anti-hamster (Abcam; ab6892). All antibodies were diluted in 10 % (v/v) normal goat serum, incubated for 30 minutes at room temperature, and visualized using 3,3-Diaminobenzine (DAB, DAKO). Sections were counterstained with Mayer's hematoxylin for 10 seconds, developed in Scott's tap water for 20 seconds, then dehydrated in ethanol and xylene. Slides were cover slipped with mounting media and scanned using the Aperio ScanScope machine (ePathology). Quantification of stained sections was performed using ImageJ.

***Opal tissue staining:*** Opal staining was carried out using the Opal staining kit (akoyabio, OP7DS2001KT) and following the below protocol; 10 mm thick sections were dewaxed in xylene and tissue hydrated in gradient ethanol washes. Antigen retrieval was performed with citrate buffer heated in a microwave pressure cooker (pH 6 for 15 minutes), followed by blocking in 10% (v/v) normal goat serum for 1 hour at room temperature. Primary antibodies were diluted in 10 % (v/v) normal goat serum and incubated for 1 hour at room temperature in a humidified chamber. Opal Polymer HRP was applied as a secondary antibody for 10 minutes at room temperature. The Opal fluorophore was then diluted in Amplification Diluent to a concentration of 1/50 and applied to the tissue for 10 minutes at room temperature. Sections were then stripped of all primary and secondary antibodies through antigen retrieval using citrate buffer heated in a microwave pressure cooker (pH 6 for 15 minutes) before the above process was repeated for each desired antibody. After the final antibody incubation, sections were incubated with spectral DAPI for 5 minutes at room temperature before mounting media was applied and sections were imaged using the Vectra imaging system. Tissue sections were then scanned using the Vectra and quantification of opal staining was performed using either InForm (Perkin Elmer) or Halo (Indica Labs).

***Preparation of single cell suspensions for flow cytometry:*** Tissues were cut into 1 mm pieces and digested in Collagenase/Dispase (Roche) and DNase I (Roche) in $Ca2^{+}$- and $Mg2^{+}$-free Hanks medium plus 5 % FCS for 30 minutes at 37°C with gentle shaking. Samples were then vortexed for 15 seconds, filtered and washed in PBS plus 5 % FCS. Single cell suspensions were blocked with FC block (Invitrogen) for 20 minutes at 4$^{o}$C, before staining with fluorophore-conjugated primary antibodies (Table S3) for 20 minutes at 4$^{o}$C in the dark. Cells were washed twice and re-suspended in PBS supplemented with 5 % FCS prior to analysis with either an Aria III cell sorter or BD FACS Canto.

Isotype antibodies (Table S3) and fluorescent-minus-one (FMO) controls were used to estimate background fluorescence in combination with either compensation beads and/or unstained controls. Dead cells were detected and excluded from analysis using Sytox Blue or Fixable Viability Dye, eF506. Tuft cells were identified as $EpCAM^{+}CD45^{-/low}CD24^{+}SiglecF^{+}$. Inflammatory ILC2s were identified as $KLRG1^{+}ST2^{-}CD90.2^{+}Lineage^{-}(CD11b^{-}CD11c^{-}CD19^{-}$Ly-

6G$^{-}$NK1.1$^{-}$CD3$^{-}$)CD45$^{+}$. Natural ILCs were identified as KLRG1$^{+}$ST2$^{+}$CD90.2$^{+}$Lineage$^{-}$(CD11b$^{-}$CD11c$^{-}$CD19$^{-}$Ly-6G$^{-}$NK1.1$^{-}$CD3$^{-}$)CD45$^{+}$. All experiments were analyzed with FlowJo software (Version 10).

**Statistical Analysis:** All experiments were conducted at least twice with ≥3 sex- and aged-matched mice per group. Where drugs were administered, animals were randomized into their corresponding treatment groups. Tumor growth was measured and recorded by an independent assessor who was blinded to the experimental conditions. No data was excluded from the analysis. Comparisons between mean values were performed with a 2-tailed Student's t-test for comparisons between 2 groups, or a one-way ANOVA for comparisons between multiple groups using Prism 10 software (GraphPad). A p value of less than 0.05 was considered statistically significant. All data is expressed as the mean ± SEM. Each 'n' or symbol represents a single mouse (biological replicate). The statistical Log-rank (Mantel-Cox) test, Hazard ratios (logrank) and median survival were calculated in Prism 10.

**Fig. S1.**

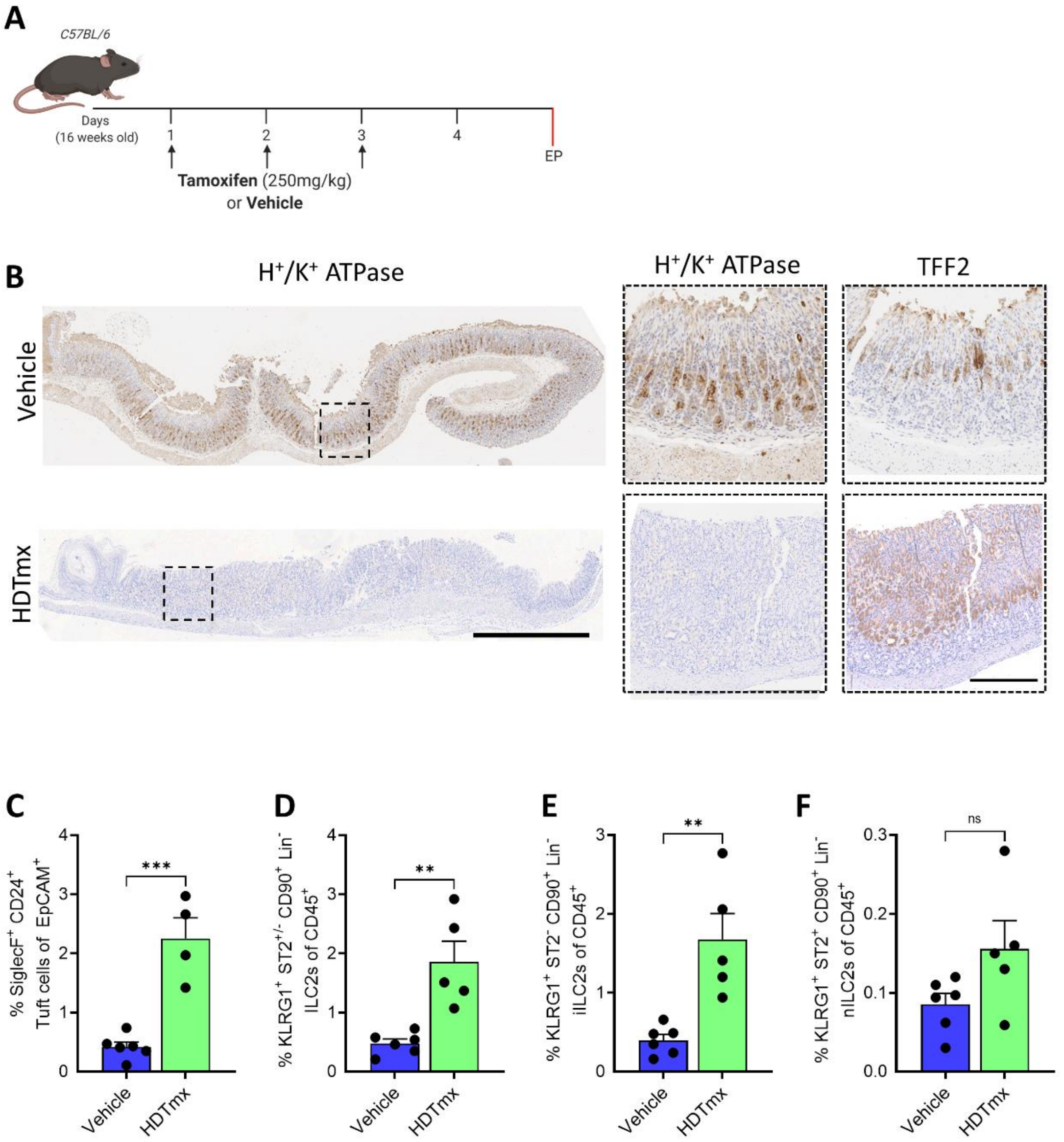
A
C57BL/6
Days
(16 weeks old)
1
2
3
4
EP
Tamoxifen (250mg/kg)
or Vehicle
B
H+/K+ ATPase
H+/K+ ATPase
TFF2
Vehicle
HDTmx
C
% SiglecF+ CD24+ Tuft cells of EpCAM+
***
Vehicle
HDTmx
D
% KLRG1+ ST2+/- CD90+ Lin- ILC2s of CD45+
**
Vehicle
HDTmx
E
% KLRG1+ ST2- CD90+ Lin- iILC2s of CD45+
**
Vehicle
HDTmx
F
% KLRG1+ ST2+ CD90+ Lin- nILC2s of CD45+
ns
Vehicle
HDTmx

**Fig. S1. Tuft cells and ILC2s are increased during Spasmolytic polypeptide-expressing metaplasia (SPEM), a precursor to gastric disease.**

(**A**) Experimental schematic to induce SPEM. 16-week-old C57BL/6 mice were treated with a high dose of tamoxifen (HDTmx, 250 mg/kg) once daily for 3 consecutive days to induce loss of parietal cells and gastric spasmolytic polypeptide-expressing metaplasia (SPEM). EP = endpoint.

(**B**) Representative image of SPEM/TFF2 and $H^+/K^+$ ATPase-stained gastric mucosa of mice as treated as described in Fig. S1A. Scale bar = 1 mm and 300 μm respectively.

(**C**) Flow-cytometry quantification of $SiglecF^+CD24^+EpCAM^+$ tuft cells in stomachs of mice as treated as described in Fig. S1A.

(**D**) Flow-cytometry quantification of total ILC2s as $KLRG1^+ST2^{+/-}CD90.2^+Lineage^-CD45^+$ in stomachs of mice as treated as described in Fig. S1A.

(**E**) Flow-cytometry quantification of iILC2s as $KLRG1^+ST2^-CD90.2^+Lineage^-CD45^+$ in stomachs of mice as treated as described in Fig. S1A.

(**F**) Flow-cytometry quantification of nILC2s as $KLRG1^+ST2^+CD90.2^+Lineage^-CD45^+$ in of mice as treated as described in Fig. S1A.

Data represents mean ± SEM, p values from Student's t-test ** $p < 0.01$, *** $p< 0.001$, n.s not significant. Each symbol represents an individual mouse.

**Fig. S2.**

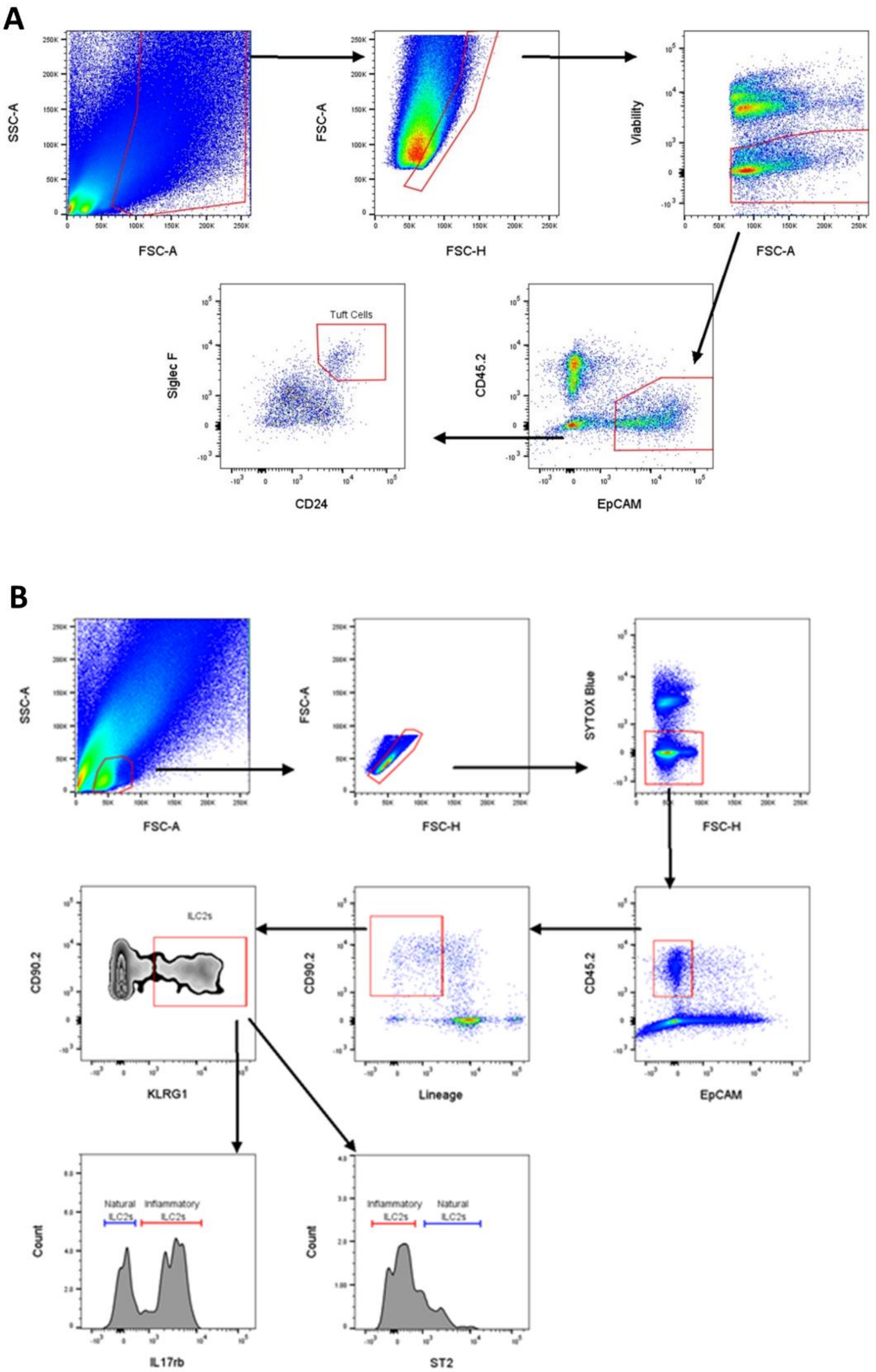
A
SSC-A
FSC-A
FSC-A
FSC-H
Viability
FSC-A
Tuft Cells
Siglec F
CD24
CD45.2
EpCAM
B
SSC-A
FSC-A
FSC-A
FSC-H
SYTOX Blue
FSC-H
ILC2s
CD90.2
KLRG1
CD90.2
Lineage
CD45.2
EpCAM
Natural ILC2s
Inflammatory ILC2s
Count
IL17rb
Inflammatory ILC2s
Natural ILC2s
Count
ST2

**Fig. S2. Gating strategy for tuft cells and ILC2s.**

(**A**) Gating strategy used to identify and sort tuft cells from mouse stomachs.

(**B**) Gating strategy used to sort ILC2s, as well as identify nILC2s and iILC2s subpopulations from mouse stomachs.

**Fig. S3.**

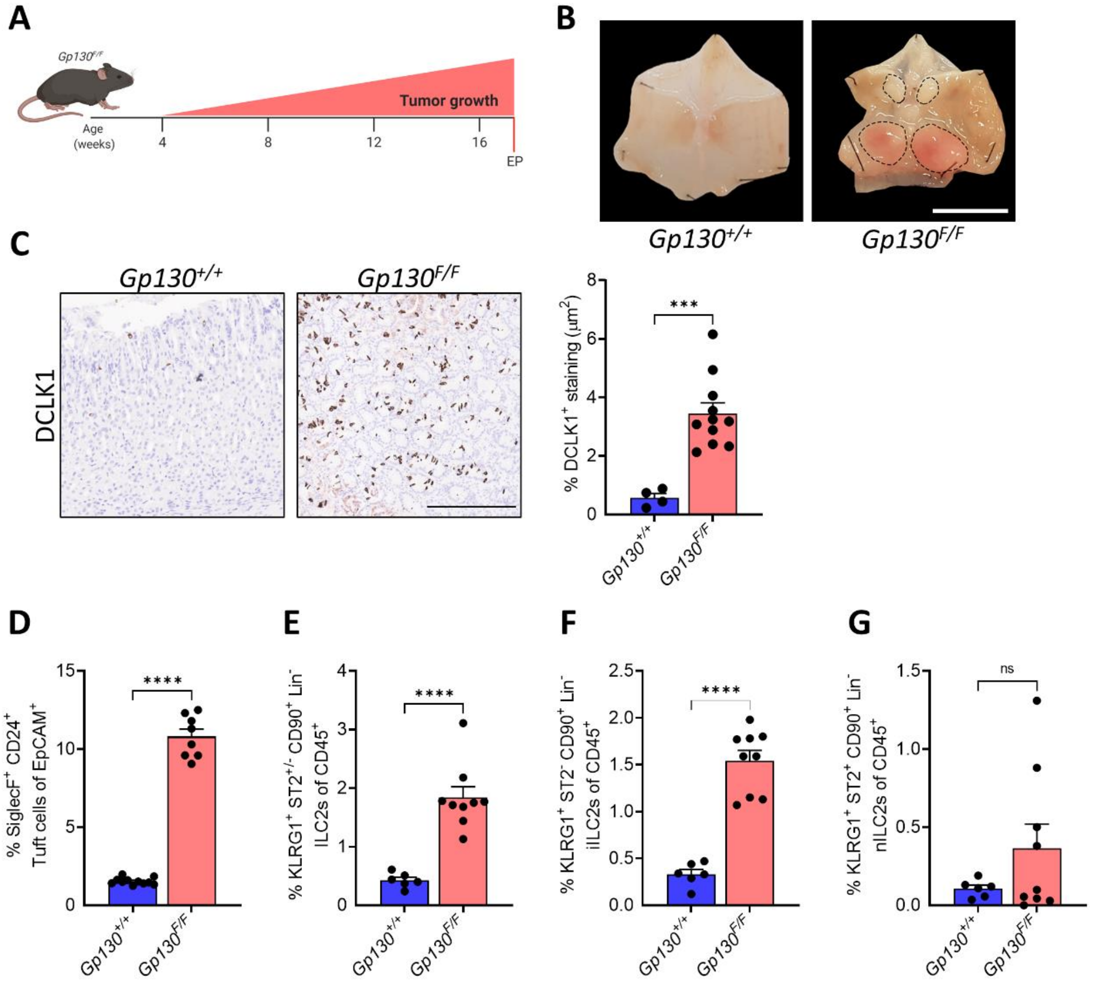
A
Gp130F/F
Tumor growth
Age (weeks)
4
8
12
16
EP
B
Gp130+/+
Gp130F/F
C
Gp130+/+
Gp130F/F
DCLK1
% DCLK1+ staining (µm2)
***
D
% SiglecF+ CD24+ Tuft cells of EpCAM+
****
E
% KLRG1+ ST2+/- CD90+ Lin- ILC2s of CD45+
****
F
% KLRG1+ ST2- CD90+ Lin- iILC2s of CD45+
****
G
% KLRG1+ ST2+ CD90+ Lin- nILC2s of CD45+
ns

**Fig. S3. Tuft cells and ILC2s are increased in during gastric tumor development.**

(**A**) Schematic of the *Gp130*$^{F/F}$ mouse model of gastric cancer, which spontaneously develops gastric adenomas from 4 weeks of age. EP = endpoint.

(**B**) Representative images of WT *Gp130*$^{+/+}$ and *Gp130*$^{F/F}$ stomachs at 17 weeks of age. Dotted circles indicate tumors, scale bar = 8 mm.

(**C**) IHC quantification of tuft cells (DCLK1$^{+}$) in *Gp130*$^{+/+}$ and *Gp130*$^{F/F}$ mice. Scale bar = 300um.

(**D**) Flow-cytometry quantification of tuft cells as SiglecF$^{+}$CD24$^{+}$EpCAM$^{+}$ cells in stomachs of *Gp130*$^{+/+}$ and *Gp130*$^{F/F}$ mice.

(**E**) Flow-cytometry quantification of total ILC2s as KLRG1$^{+}$ST2$^{+/-}$CD90.2$^{+}$Lineage$^{-}$CD45$^{+}$ cells in stomachs of *Gp130*$^{+/+}$ and *Gp130*$^{\mathrm{F/F}}$ mice.

(**F**) Flow-cytometry quantification of iILC2s as KLRG1$^{+}$ST2$^{-}$CD90.2$^{+}$Lineage$^{-}$CD45$^{+}$ cells in stomachs of *Gp130*$^{+/+}$ and *Gp130*$^{\mathrm{F/F}}$ mice.

(**G**) Flow-cytometry quantification of nILC2s as KLRG1$^{+}$ST2$^{+}$CD90.2$^{+}$Lineage$^{-}$CD45$^{+}$ cells in stomachs of *Gp130*$^{+/+}$ and *Gp130*$^{F/F}$ mice.

Data represents mean ± SEM, p values from Student's t-test *** p< 0.001, **** p< 0.0001, n.s not significant. Each symbol represents an individual mouse.

**Fig. S4.**

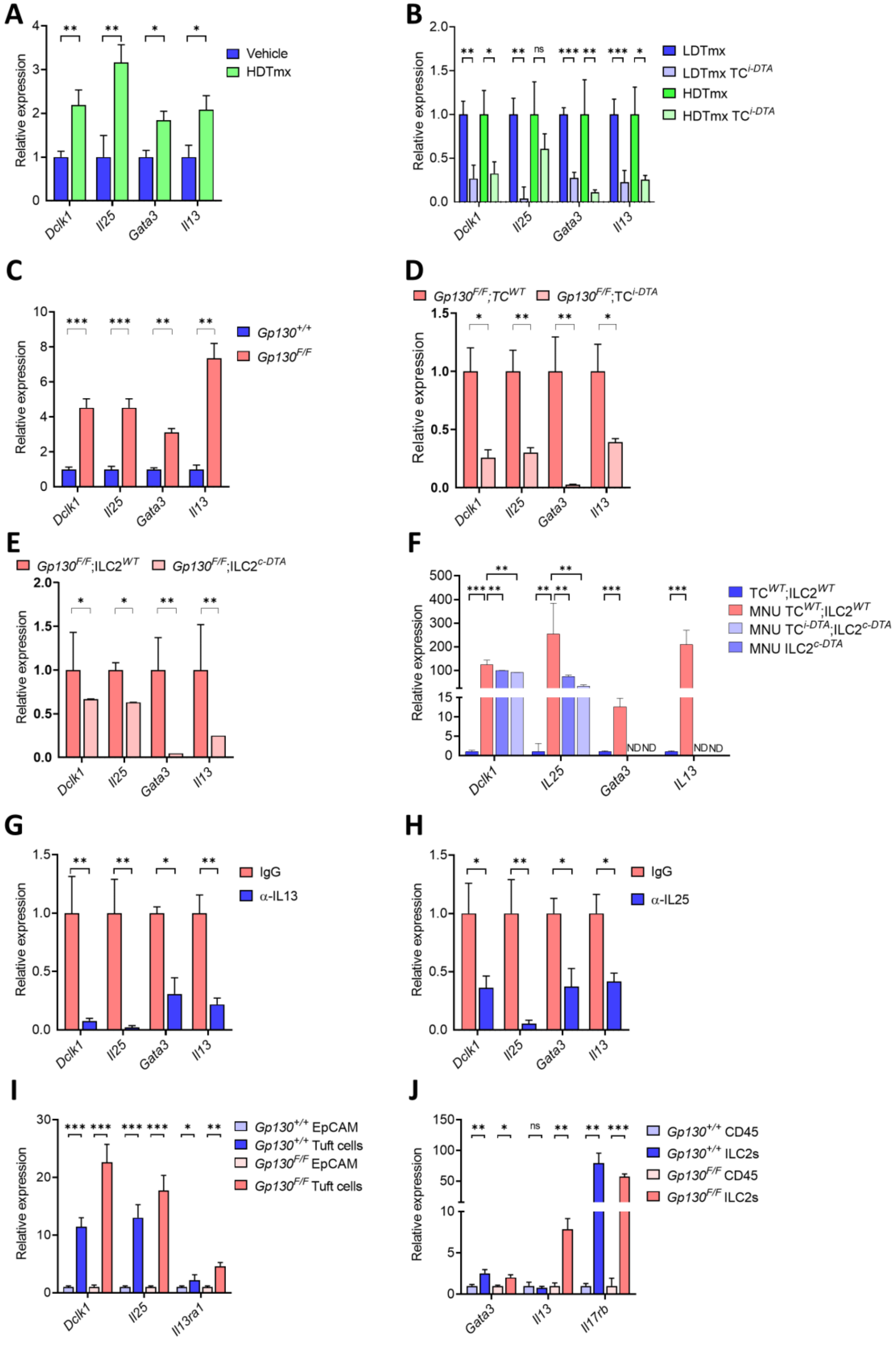
A
B
C
D
E
F
G
H
I
J
Relative expression
Vehicle
HDTmx
LDTmx
HDTmx
IgG
α-IL13
α-IL25
Dclk1
Il25
Gata3
Il13
Il13ra1
Il17rb
ND ND

**Fig. S4 Tuft cell and ILC2 markers and cytokines are increased during early gastric metaplasia and adenoma development.**

qPCR analysis for genes associated with tuft cells (*Dclk1*, *Il25)*, and ILCs2 (*Gata3*,*Il13)* in stomachs of:

(**A**) Vehicle treated and HDTmx treated mice (n=6),

(**B**) LDTmx and HDTmx treated TC$^{WT}$ and TC$^{i\text{-}DTA}$ mice (n=10).

qPCR analysis for genes associated with tuft cells (*Dclk1*, *Il25)*, and ILCs2 (*Gata3*,*Il13)* in tumors of:

(**C**) 17-week-old *Gp130*$^{+/+}$ and *Gp130*$^{F/F}$ mice (n=6),

(**D**) 17-week-old LDTmx treated *Gp130*$^{F/F}$;TC$^{WT}$ and *Gp130*$^{F/F}$;TC$^{i\text{-}DTA}$ mice (n=6),

(**E**) 17-week-old LDTmx treated *Gp130*$^{F/F}$;ILC2$^{WT}$ and *Gp130*$^{F/F}$;ILC2$^{c\text{-}DTA}$ mice (n=8),

(**F**) Age matched TC$^{WT}$;ILC2$^{WT}$, as well as MNU treated TC$^{WT}$;ILC2$^{WT}$, TC$^{i\text{-}DTA}$;ILC2$^{c\text{-}DTA}$ and ILC2$^{c\text{-}DTA}$ mice (n=6),

(**G**) IgG and α-IL13 treated *Gp130*$^{F/F}$ mice (n=4),

(**H**) IgG and α-IL25 treated *Gp130*$^{F/F}$ mice (n=4).

(**I**) qRT-PCR analysis of *Gp130*$^{+/+}$ and *Gp130*$^{F/F}$ sorted EpCAM$^{+}$ and tuft cells (SiglecF$^{+}$CD24$^{+}$) for the expression of tuft cell genes *Dclk1*, *Il25* and *Il13ra1* (n=6).

(**J**) qRT-PCR analysis of *Gp130*$^{+/+}$ and *Gp130*$^{F/F}$ sorted CD45$^{+}$ and ILC2s for the expression of *Gata3*, *Il13* and *Il17rb* (n=6).

Data represents mean ± SEM, p values from Student's t-test or ANOVA * $p < 0.05$, ** $p < 0.01$, *** $p < 0.001$, n.s not significant.

**Table S1.**

| Gene | Forward primer | Reverse primer |
|---|---|---|
| *18S* | GTAACCCGTTGAACCCCATT | CCATCCAATCGGTAGTAGCG |
| *Dclk1* | TTCAACACAGGCCCCAAG | TATCAAGAGCGGTGGTTGC |
| *Gapdh* | AAGAGGGATGCTGCCCTTA | TTTTGTCTACGGGACGAGGA |
| *Gata3* | TCGGCCATTCGTACATGGAA | GAGAGCCGTGGTGGATGGAC |
| *Il13* | CCTCTGACCCTTAAGGAGCTTAT | CGTTGCACAGGGGAGTCT |
| *Il13ra1* | TCACTTTGATGACCAACAGGAT | CAGGGGTAATTCCTCTTTACGA |
| *Il17rb* | GGACAGCCCTTCTTTGTCTG | TGCTTTTTATATTTCATTACGTGGTT |
| *Il25* | ACAGGGACTTGAATCGGGTC | TGGTAAAGTGGGACGGAGTTG |

**Table S1.** Oligonucleotide sequences for SYBR Green qPCR.

**Table S2.**

| Antibody | Reference | Manufacturer | Concentration |
| --- | --- | --- | --- |
| DCLK1 | ab31704 | Abcam | 1/1000 |
| GATA3 | SC268 | Santa Cruz Biotechnology | 1/100 |
| CD3 | MA5-14524 | Invitrogen | 1/150 |
| ChAT | AB144P | Millipore Sigma | 1/50 |
| TFF2 | Pa5-80111 | Invitrogen | 1/500 |
| $H^+/K^+$ ATPase | Ab176992 | Abcam | 1/500 |

**Table S2.** Primary antibodies for paraffin immunohistochemistry (IHC-P), Opal staining or paraffin immunofluorescence (IF).

**Table S3.**

| Antibody | Reference | Manufacturer | Conjugate | Concentration |
|---|---|---|---|---|
| CD19 | 25-0193-81 | Invitrogen | PEVio770 | 1/200 |
| CD11c | 25-0114-87 | Invitrogen | PEVio770 | 1/200 |
| CD11b | 101216 | BioLegend | PEVio770 | 1/200 |
| CD3e | 25-0031-82 | Invitrogen | PEVio770 | 1/200 |
| CD90.2 | 130-102-345 | MACS | VioBlue | 1/100 |
| KLRG1 | 138407 | BioLegend | PE | 1/200 |
| NK1.1 | 25-5941-81 | Invitrogen | PEVio770 | 1/200 |
| CD24 | 130-102-733 | MACS | APC | 1/50 |
| SiglecF | 562757 | BD Bioscience | PE-CF594 | 1/200 |
| CD45.2 | 103116 | BioLegend | APC-Cy7 | 1/200 |
| EpCAM | 11-5791-82 | Invitrogen | FITC | 1/200 |
| ST2 | 46-9335-82 | Invitrogen | PerCP-eF710 | 1/200 |
| LY6G | 560601 | BD Bioscience | PEVio770 | 1/200 |
| FC Block CD16/CD32 | 14-0161-86 | Invitrogen | | 1/100 |
| Sytox Blue | S11348 | Invitrogen | | 1/500 |
| Fixable Viability Dye | 65-0866-14 | eBioscience | eF506 | 1/1000 |
| IgG2aϰ | #400512 | BioLegend | APC | 1:200 |
| IgG2aϰ | #400230 | BioLegend | APC-CY7 | 1:200 |
| IgG2aϰ | #400208 | BioLegend | FITC | 1:200 |
| IgG2aϰ | #400908 | BioLegend | PE | 1:200 |
| IgG2aϰ | #400522 | BioLegend | PeCY7 | 1:200 |

**Table S3.** Conjugated antibodies for flow cytometry.
